# Benchmarking A Novel Quantitative PCR-based Microbiome Profiling Platform Against Sequencing-based Methods

**DOI:** 10.1101/2023.12.27.573468

**Authors:** Benjamin J. Tully, Steven E. Finkel, Christopher H. Corzett

## Abstract

**Background:** PCR-based diagnostics, predominantly utilized for pathogen detection, have faced challenges in broader microbial profiling due to disparities in genomic data availability. This study addresses this limitation by exploiting the surge in the number of microbial genomes, facilitated by advancements in next-generation sequencing (NGS) and metagenomic-assembled genomes. The primary aim was to develop and validate quantitative PCR (qPCR) assays for a wide range of gut commensals, traditionally overlooked due to inadequate genomic information. We sought to compare the efficacy of these qPCR assays against established NGS microbiome profiling methodologies - 16S amplicon and metagenomic sequencing.

**Methods:** We designed 110 species-specific qPCR assays for gut commensals using a novel proprietary *in silico* pipeline and validated the assays against stool samples from three healthy donors. The quantitative microbiome profiles were compared to taxonomic profiles generated by standard bioinformatic approaches for 16S amplicon and metagenomic sequencing. 16S amplicons were analyzed as amplicon sequence variants produced by DADA2 and metagenomic sequences were analyzed by multiple iterations of MetaPhlAn (versions 2, 3, and 4) and Kraken2/Bracken paired with two different genomic databases. The qPCR assays were assessed for their ability to detect low abundance microbes and their correlation with NGS results, focusing on taxonomic resolution and limits of quantification.

**Results:** The qPCR assays demonstrated high concordance with advanced metagenomic and the ineffectiveness of 16S amplicon methods to achieve species-level assignments. qPCR microbiome profiles were more highly correlated with the most current bioinformatic methods than the bioinformatics methods were to each other. The profile comparisons also highlight how the continued use of older bioinformatics protocols can limit results and lead to misinterpretation of data. Notably, qPCR identified taxa undetected or underestimated by metagenomic approaches, revealing limitations in current bioinformatics tools for differentiating closely related species and quantifying low abundance taxa.

**Conclusions:** This study establishes qPCR as a robust tool for large-scale microbiome profiling, offering enhanced accuracy, sensitivity, and quantitative capabilities compared to standard NGS methods. Our findings advocate for the integration of qPCR in standardizing microbiome detection, providing a pathway towards developing human microbiome profiling platforms capable of accurate species quantification. The adoption of qPCR assays could lead to more consistent, reliable, and cost-effective microbiome research and diagnostics.

## INTRODUCTION

PCR-based molecular diagnostics have become a common methodology for microbial detection over the last 40 years (Razin et al., 1984; Chen et al., 1989). However, the broadest application of these assays has been applied to the detection of microbial pathogens (Schmitz et al., 2022). While the need for accurate and rapid pathogen detection has evident clinical importance, one reason molecular-based assays have not been more readily applied to other classes of microbes has been due to the discrepancy in available high-quality, genomic information necessary to design discriminatory assays. Amongst the first sequenced genomes, pathogens now compose the largest segment of publicly available microbial genomes through the International Nucleotide Sequence Database Collaboration repositories (National Center for Biotechnology Information, European Nucleotide Archive, and DNA Data Bank of Japan) (Didelot et al., 2012). For example, the top 10 most frequently sequenced microbial organisms consist entirely of organisms with the capacity for pathogenicity, including *Escherichia coli*, *Klebsiella pneumoniae*, *Staphylococcus aureus*, and *Salmonella enterica*, and comprise 29% of all available bacterial genomes (approx. 130,000 of 395,000 genomes) (Amoutzias et al., 2022; Argimón et al., 2021; Chen et al., 2020). The investment to produce genomes for these pathogens have benefited millions of people as it has provided the necessary information about regions that are specific to those targets and disease states. Conversely, designing molecular assays for organisms without sufficient genomic information has proved to be a complicated, time-intensive, and cost-inefficient task. To compensate for the dearth of genomic information, researchers have used the 16S rRNA gene, which has extensive databases, to design molecular assays; however, the fixed length of the gene and other evolutionary constraints can make designing species-discriminating assays difficult or impossible (Fox et al., 1992; Pei et al., 2010). Instead, it would be preferable to target genomic regions suitable for molecular assays that are universally conserved exclusively within species of interest. However, without sufficient *in silico* and *in vitro* validation steps, candidate PCR-based molecular assays tend to have high (*>*80%) failure rates in real samples and environments (Bustin and Huggett, 2017; Forootan et al., 2017; Robertson and Walsh-Weller, 1998).

To address this underlying problem, we developed a proprietary approach that allows us to leverage the increased numbers of microbial genomes, made possible due to advancements in next generation sequencing (NGS) and metagenomic-assembled genomes, to identify unique genomic sites that are suitable for discriminatory qPCR assays for virtually any microbial species of interest. All assays consist of a forward and reverse primer set with an internal region suitable for a fluorogenic probe. These assays allow for the quantitative assessment of microorganisms from all environments which has previously been unavailable to the microbiome field. This technology allows for cost- and time-efficient molecular assays, suitable for methods like qPCR, dPCR (Vogelstein and Kinzler, 1999), and molecular beacons (Tyagi and Kramer, 1996), that can be readily deployed to detect microbes of interest in complex samples.

By combining species-specific assays together into a single platform, it would be possible to reliably profile large numbers of the microbial constituents of a microbiome. Currently, the most common methods of microbiome profiling leverage the short read, next generation sequencing technology (NGS) to offer deep interrogation of a single gene target, typically the 16S rRNA gene for bacteria and archaea, (amplicon sequencing) or surveying the entire genetic content of a sample (metagenomic sequencing). However, molecular detection methods offer a number of advantages over the current gold standard provided by NGS approaches (Chiu and Miller, 2019). 16S amplicon techniques have severe limitations for identifying organisms below the genus-level which subsequently makes the identification and monitoring of relevant species and/or strains difficult and can lead to misinterpretations due to a lack of taxonomic resolution (Abellan-Schneyder et al., 2021). Further, relative abundance values determined using 16S amplicon methods are confounded by the variable copy number of 16S rRNA genes amongst microbial species (Gao and Wu, 2023). Conversely, metagenomic (*i.e.*, random shotgun, *etc.*) sequencing provides high resolution taxonomy and function predictions which allows researchers to establish a deeper understanding of the microbial community. However, the inherent cost to generate and analyze data can limit the extent and scale of a research project (Bharucha et al., 2020; Greninger, 2018). Researchers must continually weigh the costs and trade-offs between deep versus shallow sequencing and/or the number of samples to include for accurate spatiotemporal resolution. Collectively, these trade-offs constrain confident and reliable limitations of detection (Forry et al., 2023). When samples have been sequenced, interpreting results requires bioinformatics expertise as well as maintaining validated and up-to-date infrastructure (Smith et al., 2022; Wright et al., 2023). These methods often produce compositional data, reflecting the relative abundance of different microbes rather than their actual quantities. Such data can lead to skewed interpretations, as changes in one microbe’s abundance affect the perceived abundance of others (Zhou et al., 2022). Additionally, these methods are prone to zero-inflated data, where low abundance microbes (*<*0.1%) are often incorrectly discarded, leading to datasets with artificially high numbers of zeros. This can significantly impact the analysis and understanding of microbial communities, as it violates the assumptions of many common statistical models, and may fail to appropriately model the true underlying relationships, leading to poor predictive performance and inaccurate conclusions (Zhang et al., 2016; Lambert, 1992). If the desired goal is to quickly monitor and enumerate known microbes in a complex sample, molecular assays provide near real-time data and feedback.

Here, we present 110 qPCR assays designed to detect commensal organisms within the human gut and deployed using a single-plex qPCR method. Historically, many of these microbes do not have qPCR assays available due to the limitations outlined above. However, here we show that these assays provide equivalent, and in some instances superior, results compared to NGS approaches, even when considering variations across several bioinformatic tools and databases. Additionally, these assays are capable of detecting microbes at much lower limits of quantification with higher degrees of certainty that is not currently achievable by NGS. We believe that the assays offer a new and unique tool to the microbiome field that will allow researchers to more effectively develop meaningful research results and, importantly, profile complex samples at scale without requiring the costs of sequencing or bioinformatic analysis.

## 1 MATERIAL AND METHODS

### 1.1 Assay Design and Validation

Branchpoint Biosciences uses the taxonomic schema established by the Genome Taxonomy Database (GTDB) to define a common understanding as to what comprises a species (Parks et al., 2021a). GTDB defines species using an predominantly automated approach with human surveillance and community feedback to identify prevailing issues and conflicts. The automated process uses relative evolutionary distance to determine delineations between higher taxonomic ranks and average nucleotide identity to establish species clusters. The schema implemented within GTDB allows for monitoring how species definitions have changed over time, while also providing links to less data rich schemas, like NCBI TaxID. Molecular assays discussed here were designed using version R207, but the underlying pipeline allows for rapid iteration when necessary (*e.g.*, changes with the recent release of version R214). For additional information on assay design and an examination of the success rate during validation of 10 molecular assays designed for 17 target species, see the **Supplemental Information**.

### 1.2 qPCR Amplification

110 species-specific primer and probe targets were identified using a proprietary bioinformatics pipeline. Primers optimized for downstream qPCR applications were synthesized (Integrated DNA Technologies, Inc., Iowa, USA) and initially assessed against at least one representative organism within the target species or by using a synthetic gBlock containing the full-length target amplicon to ensure appropriate amplification of the intended target region. PCR amplification reactions were conducted in 96-well optical plates on a Bio-Rad CFX Connect using 2*x* iTaq Universal SYBR Green Supermix (Bio-Rad, California, USA), 0.25 *µ*M of each primer, and a defined concentration of genomic DNA or gBlocks within reactions totaling 20 *µ*L. Each qPCR reaction was initiated with a 1-minute incubation at 95°C, then cycled between 15 seconds at 95°C and 30 seconds at 60°C for 40 cycles. Fluorescence intensity was detected following each cycle. Finally, a melt curve analysis of products across the range of 60°C to 95°C at 0.5°C increments was conducted. Primer sets yielding appropriate products (Ct *<*30) were next assessed against a diverse panel of 32 representative organisms observed in the human gut to ensure the absence of unintended products indicating cross-reactivity with non-target species. Primers without any signal from non-target species (C_t_ *>*35) were considered passing, whereas those resulting in unexpected products from non-target species (C_t_ *<*30) were considered failures. Some instances of inefficient amplification from unintended species (C_t_ between 30-35) were flagged for caution and additional assessment in subsequent validation experiments.

FAM-labeled fluorogenic probes were ordered (Integrated DNA Technologies) for primer sets passing cross-reactivity assessment to enable additional sequence-specific discrimination using probe-based detection. Probes were introduced at 0.25 *µ*M into qPCR reactions using 2*x* iTaq Universal Probes Supermix (Bio-Rad, California, USA) and the same amplification steps described above. Successful probe-based detection was first demonstrated using gDNA or gBlock positive controls from intended targets (C_t_ *<*30) before validated assays were assessed against diverse stool-derived templates.

Assays successfully passing initial primer positive controls, primer cross-reactivity, and probe-based detection screening steps were next validated against stool-derived communities. Using the same reaction composition and conditions for probe-based detection, 1 *µ*L (10.2-20.0 ng) of purified DNA from healthy donor stool samples spiked with known quantities of positive control organisms (described further below) was used as template material. The absolute abundance of each target organism was calculated by comparing the Ct values of each species-specific assay against that of the most abundant positive control spike-in (*P. somerae*) using the ΔC_t_ method assuming 100% reaction efficiencies. Given the known quantity of spike-in template copies within each reaction, the number of each target organism within the donor stool community could be determined.

### 1.3 Sample Collection, Sequencing, and Quality Control

Approximately 1g of stool was collected from three healthy individuals (Donor-A, -B, and -C) and resuspended in 8 mL of nitrogen-purged PBS, homogenized by shaking with glass beads, then glycerol was added to a final concentration between 15-20% before freezing at −80°C. DNA was extracted from twelve 250 *µ*L aliquots performed in parallel from each donor using the DNeasy Powersoil Pro kit (Qiagen, Hilden, Germany) according to manufacturer recommendations. Beadbeating was performed at 30 Hz for a total of 10 minutes using a TissueLyser II. Extracted DNA from each aliquot was eluted in 100 *µ*L Solution C6. All aliquots from each donor were combined and quantified using the Qubit dsDNA HS Assay Kit (Thermo Fisher, Massachusetts, USA).

One sample from each healthy donor was prepared for sequencing by GENEWIZ Azenta Life Sciences (New Jersey, USA). A total of 1,500 ng of stool-derived DNA was used to construct each sequencing sample. As a positive control, the gDNA for *Porphyromonas somerae*, *P. uenonis*, and *P. asaccharolytica* were spiked into each sample at relative concentrations of 1.0%, 0.1%, and 0.001%, respectively. All three samples were screened for the presence of the spike-ins using previously validated qPCR assays. Samples underwent 16S (V3-V4) amplicon and metagenomic sequencing to 200k and 300M reads per sample, respectively.

Metagenomic sequences were screened for quality using fastp (v.0.23.4) (Chen et al., 2018) with a sliding window mean quality filter of Q28, minimum length of 75 bp, and adapter screening (R1 = AGATCGGAAGAGCACACGTCTGAACTCCAGTCA and R2 = AGATCGGAAGAGCGTCGTG-TAGGGAAAGAGTGT) from the 5’ and 3’ end of the forward and reverse reads. Further, any reads that were mapped to the human genome (GRCh38.p14) were removed using minimap2 (v.2.24-r1122; parameters: -ax sr) (Li, 2018) and samtools (v.1.9) (Li et al., 2009). 16S amplicon sequences were screened for quality during implementation of the “denoise-paired” function of DADA2 (Callahan et al., 2016) within QIIME2 (v.qiime2-2022.11) (Bolyen et al., 2019).

### 1.4 Bioinformatic Analysis

#### 1.4.1 16S amplicon

The representative amplicon sequence variant (ASV) sequences produced from the DADA2 “denoise-function” were classified with the subcommand “feature-classifier classify-sklearn” which applied a pre-trained Naive Bayes classifier built using the Silva R138 99% operational taxonomic units (OTUs) full-length available through QIIME2Docs (docs.qiime2.org/; “Data Resources”; MD5: b8609f23e9b-17bd4a1321a8971303310). The tab-delimited taxonomy file was merged with the sample metadata to create a “biom” format with the biom subcommand “add-metadata” and converted into tab-delimited file using the subcommand “convert.” For analysis purposes, the relative abundance of ASVs with the same species designation were combined. Due to a low number of species assignments, the 16S amplicon results were not included in the statistical analysis approach described below.

#### 1.4.2 Metagenomic

Taxon tables were generated for all three samples using multiple bioinformatic tools and variations of available databases (described below). Analyses were performed on two versions of the data: (1) All reads remaining after quality control and removal of host signal; and (2) subsampled set of paired reads used to simulate a typical “shallow” metagenomic dataset (10M paired reads). Subsampling was performed using seqtk (v.1.3-r106; https://github.com/lh3/seqtk). Samples were analyzed using MetaPhlAn2 (v.2.7.7) (Truong et al., 2015), MetaPhlAn3 (v.3.1.0) (Beghini et al., 2021), MetaPhlAn4 (v.4.0.3) (Blanco-Míguez et al., 2023), and Kraken2/Bracken implemented with the GTDB (v.R207) and Standard (v.12/09/2022) databases. The three versions of MetaPhlAn were implemented using their corresponding default settings with the “analysis type” flag set to “rel ab w read stats.” Taxon table results produced by MetaPhlAn4 were converted to the GTDB taxonomy schema using the script sgb to gtdb profile.py available through the Utilities directory hosted on the MetaPhlAn4 GitHub page (https://github. com/biobakery/MetaPhlAn). Kraken2 (v.2.1.2) was implemented using both databases with confidence cutoff set to 0.1 and the “report-minimizer-data” flag (Wood et al., 2019). Bracken (v.2.8) was run on the requisite Kraken2 reports with a read distance of 150 and set to the species level (parameters: -r 150 -l S) (Lu et al., 2017).

#### 1.4.3 Artificial Metagenomes

A list of 24 bacterial species was constructed informed by the taxa that were identified by at least one metagenomic bioinformatic analysis and not captured by the qPCR assays (see Statistical Analyisis and Outliers) and phylogenetic neighbors. For example, as with Donor-A, the metagenomic bioinformatic analysis predicted the presence of *Faecalibacterium prausnitzii E*, while the corresponding qPCR assay was negative. Thus *F. prausnitzii E* and all related *F. prausnitzii* groups, defined by GTDB, were added to the list. The list was used to construct artificial metagenomes. A random subsample of 11 species was selected from the list of 24. At random, two or three species were assigned relative abundance from 0 *−* 0.01 and all remaining species were assigned relative abundance from 0.01 *−* 0.15, totaling to a sum of 1.0. From these settings, the Metagenomic Sequence Simulator (MeSS; https://github.com/metagenlab/MeSS) was provided the GTDB type genome NCBI Assembly ID and produced simulated metagenomes approximating a 5M paired-end Illumina HiSeq with 150bp read lengths (replication random seed = 78). The paired-end reads were processed with Kraken2/Bracken implemented with the GTDB (v.R207), as described above. Taxa detected at *≥*0.04% relative abundance were plotted for comparison against the defined input.

### 1.5 Statistical Analysis and Outliers

To determine how the various molecular and bioinformatics methods interpreted the taxa data, we screened the results of all methods for 110 target taxa. Any taxa that were reliably quantifiable in at least one of the data sets, including bioinformatics methods, were retained for statistical analysis. “Reliably quantifiable” was defined as either *≥*1 copy per reaction for a qPCR assay or the relative abundance value associated with a 10% false positive rate as determined previously (Bracken 0.04%, MetaPhlAn2/3/4 0.01%) (Parks et al., 2021b). The total signal of all taxa of interest was calculated from the sum of values determined for each individual target. This was used to determine the “relative signal contribution” for each taxon by dividing the signal from one target taxon by the sum of the total signal. Relative signal abundance was compared in a pairwise manner between all methods using linear regression analysis and analysis of variance (ANOVA) using the “statsmodels” Python library (v.014.0). Residuals were determined by comparing each taxon to the linear regression model and were used to calculate the corresponding standardized residuals. A standardized residual *≥*3 was used to identify statistical outliers between methods.

Outliers were assessed in a number of ways with the goal of determining if there was an underlying reason for the discrepancy. In many instances, this involved comparing the GTDB species assignment with the NCBI TaxID and/or species-level genome bins (SGBs) in a corresponding MetaPhlAn database. For MetaPhlAn4, publicly available SGBs could be assessed with the addition of a GTDB taxonomy as determined by the GTDB-Tk (v.2.1.1; R207) (Chaumeil et al., 2022) subcommand “classify wf.” Further, if necessary, a phylogenetic tree was constructed using GToTree (v.1.6.35) by downloading the corresponding “type genomes” as assigned by GTDB from NCBI and selecting an appropriate set of single copy marker genes (*e.g.*, *Faecalibacterium prausnitzii* used the *Firmicutes* single copy gene set) (Lee, 2019). Output phylogenetic trees were visualized using the interactive Tree of Life (iToL; https://itol.embl.de/).

## 2 RESULTS

Throughout the results and discussion, when suitable, quantities will be presented as a range of values obtained from all three samples.

### 2.1 qPCR

Based on the “reliably quantifiable” thresholds, 106 target taxa were identified across the three donor samples with 81 detected by qPCR (Supplemental Data 1). Using *P. somerae* as a single point calibration value for the qPCR results, the estimated total copy number of detected microbes in each sample was 1.7 *−* 2.9 *×* 10^6^ per reaction. Each reaction represents approximately *≈*1.4-3.0 mg of processed fecal material. With a C_t_*_<_* _35_ threshold, *Gemella morbillorum* had the lowest detectable copy number for Sample the qPCR reactions with *≈*7 copies detected in the sample from Donor-A. An additional 17 reactions exceeded C_t_*_<_* _35_ and corresponds to 1 *−* 4 copies per reaction. *Coprococcus eutactus A* had the highest copy number with 337,600 copies detected in the sample from Donor-B. The mean number of copies detected per taxa in each sample was 2.2 *−* 3.8 *×* 10^5^ copies (median: 28 *−* 3, 935 copies).

### 2.2 Metagenomes

For the three metagenomic samples, we received 236-296M raw reads (35-44 Gbp). Between 0.9-1.1% of the reads were removed during the quality control step and another 0.1-0.4% of reads were removed when recruited against the human genome (232-292M QCed reads). The Bracken approaches using the GTDB and Standard databases classified 81-84% and 42-51% of reads, respectively (Supplemental Table 1). Each bioinformatic approach produced a variable number of species-level assignments above their respective “reliably quantifiable” threshold of a 10% false positive rate (MetaPhlAn2 74-79 identified species; Bracken-GTDB 193-245 identified species) (Supplemental Table 2). Collectively, taxa detected by qPCR account for 31-41% of the total relative abundance determined as part of the MetaPhlAn4 profile. For each sample, it was determined which taxa were detected by qPCR but fell below the “reliably quantifiable” threshold for all five bioinformatic approaches (Supplemental Data 2). When using the full dataset (300M reads), 12-16 taxa were below the bioinformatic-based “reliably quantifiable” detection limit per sample (Table 1) with an additional 1-2 taxa undetectable when using the subsampled dataset (10M reads). Conversely, the bioinformatic methods detected between 7-12 unique taxa that were reported as absent via qPCR (Table 1; Supplemental Data 2). Some of the methods reported the same taxon as present. None of the “bioinformatic-only” taxa were reported by MetaPhlAn4 and the Bracken methods, paired with both the GTDB and Standard databases, returned a majority of these results (GTDB DB: 5-8 nonunique taxa; Standard DB: 2-4 nonunique taxa).

**Table 1.**
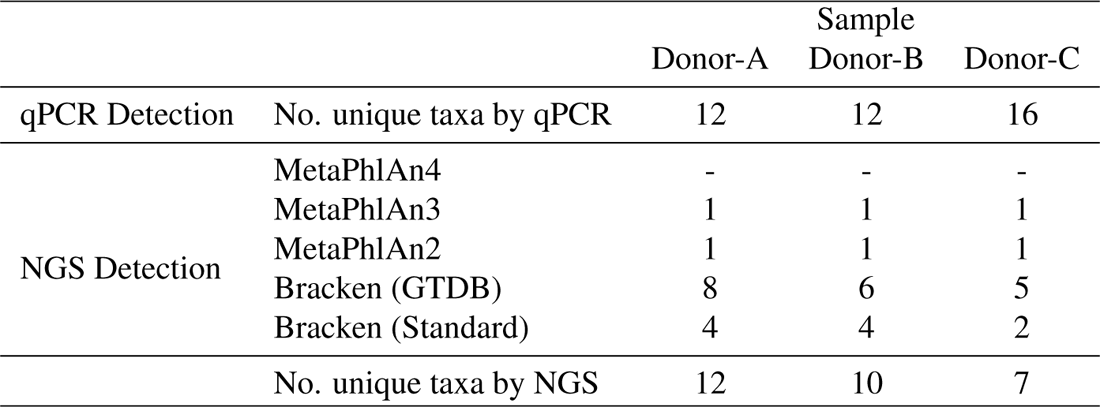
Differentially detected taxa.

### 2.3 16S Amplicon

The raw dataset of 16S amplicons consisted of 234k-377k paired-end reads. The DADA2 denoising step removed a vast majority of sequences due to detected chimerism (88-90% reads removed) retaining 25k-44k reads per sample. The DADA2 analysis generated 923 ASVs (Supplemental Data 3). Only 292 ASVs had assignments to the species-level (31.6%). When considering ASVs that provide binomial nomenclature, excluding non-specific species identifiers or placeholder names (*e.g.*, *Alistipes sp.* or uncultured bacterium, respectively), only 74 ASVs (8.0%), ultimately assigned to 31 different species, remained in the dataset. The overall lack of species-level detection prevented inclusion of the 16S amplicon data in the subsequent statistical analyses. Taxa with species-level identification were detected at 0.01-2.27% relative abundance.

### 2.4 Statistical Analysis

The qPCR assays, MetaPhlAn4, and Bracken-GTDB all share an underlying taxonomy structure, while the other methods rely on databases derived from other sources. MetaPhlAn2/3 uses reference genomes from NCBI exclusively. In the pairwise comparison, approaches that share overlapping taxonomy structure generally have higher coefficients of determination (*R*^2^) and lower sum of squares (SS) (Table 2). However, each sample has differences. For the samples from Donor-B and -C, the qPCR results have a higher coefficient of determination (*i.e.*, better fitting linear model) when compared to MetaPhlAn4 and Bracken-GTDB (*R*^2^ *≥* 0.79) than when comparing MetaPhlAn4 and Bracken-GTDB (*R*^2^: 0.72 *−* 0.73). In Donor-A, qPCR and MetaPhlAn4 are highly correlated (*R*^2^ = 0.89) and have better fit than MetaPhlAn4 and Bracken-GTDB (*R*^2^ = 0.78), but qPCR and Bracken-GTDB have a lower coefficient of determination (*R*^2^= 0.74). The highest coefficients of determination tended to be between the bioinformatic methods whose databases and taxonomies are derived using the same taxonomic structure and databases. Comparisons including Bracken-Standard tended to have the lowest coefficients of determination (*R*^2^ = 0.13 *−* 0.77).

**Table 2.** Pairwise Regression Analysis Between Methodologies.

| Comparison |  | Donor-A |  | Donor-B |  | Donor-C |  |
| --- | --- | --- | --- | --- | --- | --- | --- |
| | | $R^2$ | SS | $R^2$ | SS | $R^2$ | SS |
| qPCR | MetaPhlAn4 | **0.90** | **70.84** | **0.91** | **67.64** | **0.83** | **70.77** |
|  | MetaPhlAn3 | 0.56 | 379.42 | 0.72 | 284.08 | 0.32 | 495.29 |
|  | MetaPhlAn2 | 0.32 | 934.232 | 0.43 | 692.55 | 0.28 | 582.75 |
|  | Bracken (GTDB) | **0.75** | **111.21** | **0.80** | **122.29** | **0.80** | **68.51** |
|  | Bracken (Standard) | 0.47 | 429.36 | 0.63 | 377.05 | 0.32 | 510.43 |
| MetaPhlAn4 | MetaPhlAn3 | 0.64 | 309.52 | 0.60 | 405.33 | 0.45 | 400.65 |
|  | MetaPhlAn2 | 0.50 | 683.14 | 0.60 | 482.67 | 0.41 | 477.84 |
|  | Bracken (GTDB) | **0.79** | **91.83** | 0.73 | 167.96 | **0.75** | **87.52** |
|  | Bracken (Standard) | 0.33 | 548.21 | 0.51 | 492.20 | 0.38 | 467.68 |
| MetaPhlAn3 | MetaPhlAn2 | **0.80** | 268.41 | 0.42 | 713.44 | **0.94** | **51.52** |
|  | Bracken (GTDB) | 0.59 | 180.79 | 0.58 | 263.77 | 0.27 | 254.26 |
|  | Bracken (Standard) | 0.40 | 491.51 | **0.78** | 222.03 | 0.67 | 245.97 |
| MetaPhlAn2 | Bracken (GTDB) | 0.34 | 287.97 | 0.19 | 503.28 | 0.21 | 274.05 |
|  | Bracken (Standard) | 0.14 | 701.95 | 0.37 | 633.94 | 0.49 | 385.01 |
| Bracken (GTDB) | Bracken (Standard) | 0.41 | 482.41 | 0.48 | 520.97 | 0.32 | 514.12 |

$R^2$ : coefficient of determination
SS: ANOVA Sum of squares
**Bold** indicates $R^2 \geq 0.75$ or $SS \leq 150$

### 2.5 Outlier Microbes

Using standardized residuals (*≥*3), 18 taxa were identified as statistical outliers for at least 1 method comparison (Supplemental Data 4). Three taxa (*Agathobacter rectalis* [formerly *Eubacterium rectale*], *Faecalibacterium prausnitzii*, and *Phocaeicola vulgatus* [formerly *Bacteroides vulgatus*]) were identified as outliers for qPCR in all three samples. It should be noted that *P. vulgatus* was only considered an outlier between qPCR versus MetaPhlAn2 and Bracken-Standard. Nine taxa were identified only as outliers in one sample. Amongst these, 2 taxa (*Bifidobacterium longum* and *Blautia A obeum*) was identified as an outlier comparing amongst bioinformatic methods and not in comparison to the qPCR result. In no instance was a taxon determined to be an outlier for all bioinformatic methods in comparison to qPCR. Additionally, for instances where a taxon was considered an outlier for either MetaPhlAn4 or Bracken-GTDB, the other methodology did not report the taxon as an outlier.

### 2.6 Artificial Metagenomes

The artificial metagenomes were seeded from 24 species that were either detected or closely phylogenetically related to the taxa reported by “bioinformatic-only” methods (Supplemental Data 5). Each 11-species artificial metagenome was processed with Bracken using the GTDB (v.R207) database. For the taxa that were included as the source genomic material for the artificial metagnomes, the expected and Bracken-determined relative abundance values were in general agreement (mean = 2.54% *±* 3.15). However, Bracken-GTDB reported between 2-24 additional taxa that were not present in the source artificial metagenome (above the 0.04% reliably quantifiable cutoff) which ranged in relative abundance from 0.04-3.44% (mean = 0.23% *±* 0.43). In four of the five artificial metagenomes, Bracken-GTDB reported between 3-6 distinct Faecalibacterium-related species (*F. prausnitzii*, *F. prausnitzii A*, *F. prausnitzii D*, etc.) and returned 4-16 Faecalibacterium-related species, including species groups with placeholder identifiers (*e.g.*, *Faecalibacterium sp900539945*). Placeholder identifiers were common in the additional taxa returned by Bracken-GTDB.

## 3 DISCUSSION

### 3.1 Variability amongst the positive controls

Three type strains of *Porphyromonas* (*P. somerae*, *P. uenonis*, and *P. asaccharolytica*) were used as positive controls amended to each donor sample at 1%, 0.1%, and 0.001%, respectively, of the total extract DNA (Table 3). In examining the bioinformatic approaches used to analyze both NGS methods, how different tools “detected” these three taxa reveal some of the many pitfalls inherent in microbiome profiling (which will also be explored further below). For the metagenomic approaches, several species were not present in certain databases: *P. somerae* was not present in MetaPhlAn2 or MetaPhlAn3 and *P. uenonis* was not present in MetaPhlAn4 or Bracken-Standard. None of the other methods were entirely accurate in determining the relative abundance of *P. somerae* (amended at 1.0% total DNA; mean values across all donors: MetaPlAn4: 0.912%, Bracken-GTDB: 1.153%, Bracken-Standard: 0.950%). Bracken-GTDB overestimates *P. somerae* relative abundance, but both MetaPhlAn4 and Bracken-Standard are approximately correct, especially when considering 0.1-0.4% of human gDNA was removed prior assessing the sequencing results. *P. asaccharolytica* (amended at 0.001% total DNA) was detected by all methods, but at relative abundance values well below the “reliably quantifiable” threshold for all of the bioinformatic methods and for three methods at least one order of magnitude different (mean values across all donors: MetaPhlAn4: 0.0001%, MetaPhlAn3: 0.0020%, MetaPhlAn2: 0.0119%, Bracken-GTDB: 0.0034%, Bracken-Standard: 0.0290%). On a practical level, a method that overestimates the abundance of a taxon by an order of magnitude impacts reliability, especially if there is a need to monitor “dose responses” (*i.e.*, where a taxon functions as a biomarker at a specific threshold). When the method does report an accurate value, the problem becomes identifying these results amongst an increasing number of false positives (Parks et al., 2021b). The practice of discarding taxa below a set threshold is commonly used. A conservative approach to avoid inclusion of false positives, such practices have an impact on statistical analyses due to artifacts like zero-inflation, caused by the artificially high numbers of zeros, and can lead to misinterpretation of the data or superfluous conclusions about taxa abundance and presence Zhang et al. (2016). As the monitoring of the gut microbiome is almost exclusively performed using stool samples and microbial signals are affected by the physical and chemical process of stool formation in the proximal colon, signals originating from upper and lower intestinal tracts may be regularly overlooked by sequencing methods with poor limits of detection (Mukhopadhya et al., 2022).

**Table 3.** *Porphyromonas* spike-in results across methods.

|  | Target Abund. | Spike-in |  |  |
| --- | --- | --- | --- | --- |
|  |  | *P. somerae*<br>1.0% | *P. uenonis*<br>0.1% | *P. asaccharolytica*<br>0.001% |
| Donor-A | qPCR | 40,098 | 2,921 | 10 |
|  | 16S (%) | 1.57 | 0.13 | **n.d.** |
|  | Metagenomic (%) | 0.94 | **n.d.** | **n.d.** |
| Donor-B | qPCR | 78,624 | 5,166 | 13 |
|  | 16S (%) | 1.39 | **n.d.** | **n.d.** |
|  | Metagenomic (%) | 0.93 | **n.d.** | **n.d.** |
| Donor-C | qPCR | 78,624 | 5,511 | 7 |
|  | 16S (%) | 0.76 | 0.19 | **n.d.** |
|  | Metagenomic (%) | 0.87 | **n.d.** | 0.0003 |

*P. somerae* used as single point calibration
n.d.: not detected
"Metagenomic" reports MetaPhlAn4 percent relative abundance results

**Table 4.** Quantitative Detection of Taxa Below Sequencing “Reliably Quantifiable” Threshold.

| Target Taxa | Donor-A<br>(genome copies per reaction) | Donor-B | Donor-C |
| --- | --- | --- | --- |
| *Anaerococcus obesiensis* | **4** | **75** | **54** |
| *Anaerococcus vaginalis_C* | 0 | **48** | 0 |
| *Bifidobacterium longum* | 187,822 | **8** | 5,594 |
| *Bifidobacterium pseudocatenulatum* | 185,480 | **3** | **6,211** |
| *Blautia_A obeum_B* | **7** | **39** | 591 |
| *Bulleidia moorei* | 0 | 0 | **278** |
| *Clostridium_Q symbiosum* | 0 | **1** | **9** |
| *Enterocloster asparagiformis* | **81** | **53** | **102** |
| *Fusobacterium nucleatum* | 0 | 0 | **1** |
| *Fusobacterium vincentii* | 0 | 0 | **2** |
| *Gemella morbillorum* | **7** | **18** | **126** |
| *Lancefieldella parvula* | **1** | 0 | **1** |
| *Parvimonas micra* | **1** | **2** | 1,197 |
| *Pauljensenia odontolytica_A* | **1** | **10** | 0 |
| *Pauljensenia odontolytica_B* | **16** | 0 | **64** |
| *Pauljensenia odontolytica_C* | 0 | **3** | 0 |
| *Peptostreptococcus anaerobius* | **3** | 0 | **18** |
| *Peptostreptococcus stomatis* | 0 | 0 | **321** |
| *Porphyromonas asaccharolytica* | **10** | 13 | **7** |
| *Streptococcus parasanguinis_E* | **1,365** | 0 | **893** |
| *Streptococcus parasanguinis_F* | **1,196** | 0 | **4,135** |
| *Streptococcus salivarius* | 10,520 | **99** | 2,738 |

**Bold** taxa detected by qPCR below the NGS "reliably quantifiable" threshold

The 16S amplicon results also failed to capture the positive controls consistently (Table 3). *P. asaccharolytica* was below the detection limit in all three samples, *P. uenonis* (amended at 0.1% total DNA) was undetected in Donor-B, and *P. somerae* was simultaneously over- and underestimated across the samples (Donor-A: 1.53%, Donor-B: 1.39%, Donor-C: 0.76%). Further, these relative abundance values had to be determined by combining 16S-based species assignments for multiple different ASVs (*P. somerae*: 8 ASVs, *P. uenonis*: 3 ASVs), none of which overlapped across samples (Supplemental Data 3). Based on validated qPCR assays, none of the donor samples contain any of the three *Porphyromonas* species. As such, we would expect that each species should have only one detectable ASV and the same spike-in ASVs should be observable across all three samples. The 16S amplicon method could not detect the low abundance spike-in, which, as highlighted above, would impact interpretations for microbes rare within stool samples, and severely overinflated the diversity of the other positive controls. These inherent discrepancies impact basic ecological measurements such as alpha- and beta-diversity.

### 3.2 Lack of resolution amongst 16S amplicon results

The lack of species resolution by 16S amplicon approaches is a known issue for the methodology. In this dataset, less than 10% of the representative sequences could be assigned a species designation. Amongst those with species annotations, only seven had overlap with the taxa queried using the qPCR assays. The approach used to compare the qPCR results with the metagenomic bioinformatic results (see above, relative signal contribution) is not available for the 16S amplicon results due to low number of overlapping taxa. Due to the high correlation between qPCR and MetaPhlAn4 (see below), we will compare 16S amplicon relative abundance with the values determined by MetaPhlAn4 (Supplemental Data 3). For the seven taxa across three samples (21 total comparisons), 16S and MetaPhlAn4 are in agreement twice (10%); correctly not detecting *Phocaeicola dorei* and detecting *Bacteroides uniformis* (1.07% relative abundance) in Donor-C. For 11 taxa 16S underestimated abundance (mean: *−*1.54%, range: 0.001-4.93%) and overestimated for the remaining eight (mean: +0.75%, range: 0.30-1.50%). Again, we highlight this well-known methodological issue here because of the impact these types of results would have on accurately interpreting microbiome profile data. This impact is muted when using 16S amplicons to broadly compare samples and providing a methodology to compare differences of large-scale changes across samples. However, the more important the combination of exact microbe identification and accurate numerical representation are for microbiome profile interpretation, the less utility 16S amplicons possess. This is especially important to consider for efforts that aim to use 16S amplicon sequencing in a diagnostic capacity or monitor microbiome profiles in response to therapeutic interventions.

The donor samples were screened for 110 species-level taxa using Branchpoint Biosciences proprietary qPCR assays. Of these, 106 were detected by either qPCR and/or at least one metagenomic analysis (Figure 1). A handful of taxa (n = 18) were identified as statistical outliers in at least one pairwise comparison of the methods.

**Figure 1.**
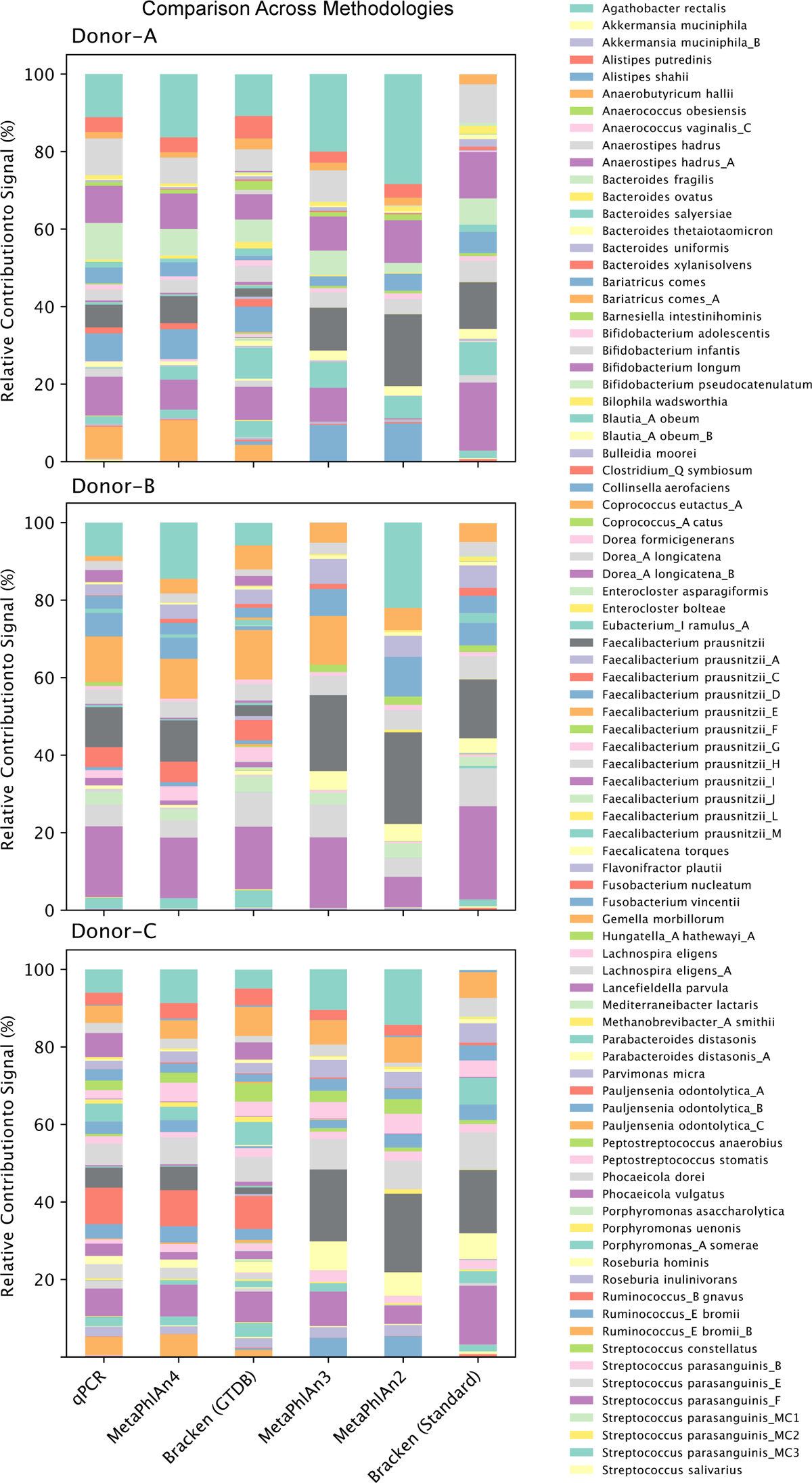
Comparison of relative contribution to signal for all three donors. qPCR assays are directly compared to five bioinformatic methods.

### 3.3 High degrees of concordance between qPCR and the latest metagenomic approaches

In general, the qPCR assays have a higher degree of correlation with each of the methods than the two bioinformatic methods have with each other (Table 2). As noted above, the qPCR assays, MetaPhlAn4, and Bracken-GTDB share a common underlying taxonomic structure, based on the genome taxonomy methodology implemented by the GTDB. This demonstrates that despite the similarity in methodological approach between these two metagenomic methods - both MetaPhlAn4 and Bracken recruit reads to a fixed database using similar genome-based methods to define the species groups - there is an inherent level of incongruence that arises due to how the specific methodologies have automated defining those species groups and manifest in the taxonomic profiles. As the qPCR assays are predefined and undergo a degree of manual curation during assay design and validation, they can be designed to account for discrepancies that arise during these automated processes. As a result, the qPCR assays closely reflect the bioinformatic methods they share a taxonomic schema with compared to the methods using a different schema. Additionally, the qPCR assays can describe species groups that more precisely reflect meaningful biological and evolutionary distinctions. As a specific example, the qPCR result for *Collinsella aerofaciens* is identified in the Donor-B sample as an outlier for Bracken-GTDB, but not MetaPhlAn4, with relative contribution values of 6.03%, 0.99%, and 5.46%, respectively. In this instance, *C. aerofaciens* is defined as a single species group by MetaPhlAn4, but has been split into at least 32 different species by GTDB, many with only a single genome representative. The 157,218 reads that recruit to *Collinsella* genus by the Bracken-GTDB method for this sample are split across 395 species-level taxa, with top assignment to a placeholder designation “*Collinsella sp905216145*.” Manual inspection of the *C. aerofaciens* phylogenetic tree revealed that a single encompassing species definition was appropriate (Supplemental Figure S2). Clarifying this discrepancy helps to ensure that our qPCR method provides accurate microbiome results that can provide consistent, reliable quantification and profiling.

Methods that do not share an underlying taxonomic structure have lower coefficients of determination (*R*^2^). The lower correlation values are also impacted by the incongruencies in the underlying reference databases. Both MetaPhlAn2 and MetaPhlAn3 were constructed when fewer reference genomes were available (7,500 and 13,500 species, respectively, compared to 26,970 species in MetaPhlAn4). This is reflected in the total number of taxa detected by each method and accounts for the decreasing correlation between methods as the time between initial release increases. The impact of different scales and contents of the databases on taxonomy results can be seen in the comparison between the GTDB and Standard databases used in conjunction with Kraken/Bracken. The GTDB database contains reference information for 85,000+ microbial genomes, while the Standard database contains 21,000+ microbial, viral, and eukaryotic genomes from NCBI RefSeq. Consequently, Bracken paired with the GTDB database assigns approximately 30% more reads to microbial taxa than when paired with the Standard database and detects approximately twice as many taxa.

### 3.4 Limitations of the underlying database structure

The detection of *F. prausnitzii* by qPCR is consistently in agreement with MetaPhlAn4 and not Bracken-GTDB and MetaPhlAn2/3. This discordance is a direct result of the underlying database structure. Both MetaPhlAn2 and MetaPhlAn3 are built on older database structures (published in 2015 and 2021, respectively) and use the canonical species definition of *F. prausnitzii*. It has since been recognized that the encompassing definition of *F. prausnitzii* is inaccurate and that multiple species groups are defined within that phylogenetic space (Zou et al., 2021; Sakamoto et al., 2022; Filippis et al., 2020). Our qPCR assays, MetaPhlAn4, and Bracken-GTDB define multiple related *F. prausnitzii* species. However, similar to *C. aerofaciens* and as demonstrated with the artificial metagenome results, the Bracken-GTDB method distributes the *F. prausnitzii* signal across related species (Figure 2). For example, in Donor-A the qPCR and MetaPhlAn4 detected the species *F. prausnitzii*, *F. prausnitzii C*, *F. prausnitzii D*, and *F. prausnitzii G* and observe different combinations of *F. prausnitzii* species in the other samples. However, in all three samples, Bracken-GTDB distributes reads across multiple *F. prausnitzii* species. This is the result of how Bracken assigns reads through the process of matching k-mers across entire genomes. The close phylogenetic relationship between these species results in reads matching non-discriminatory segments of competing genomes and produces a signal that generally underestimates species that are present and overestimates species that are absent. MetaPhlAn4 identifies unique discriminatory markers for each species and maps directly to those genes, avoiding non-specific read mapping. To be considered fully validated, the qPCR primers must also be discriminatory and exclusive to the target species, avoiding issues of cross species identification.

**Figure 2.**
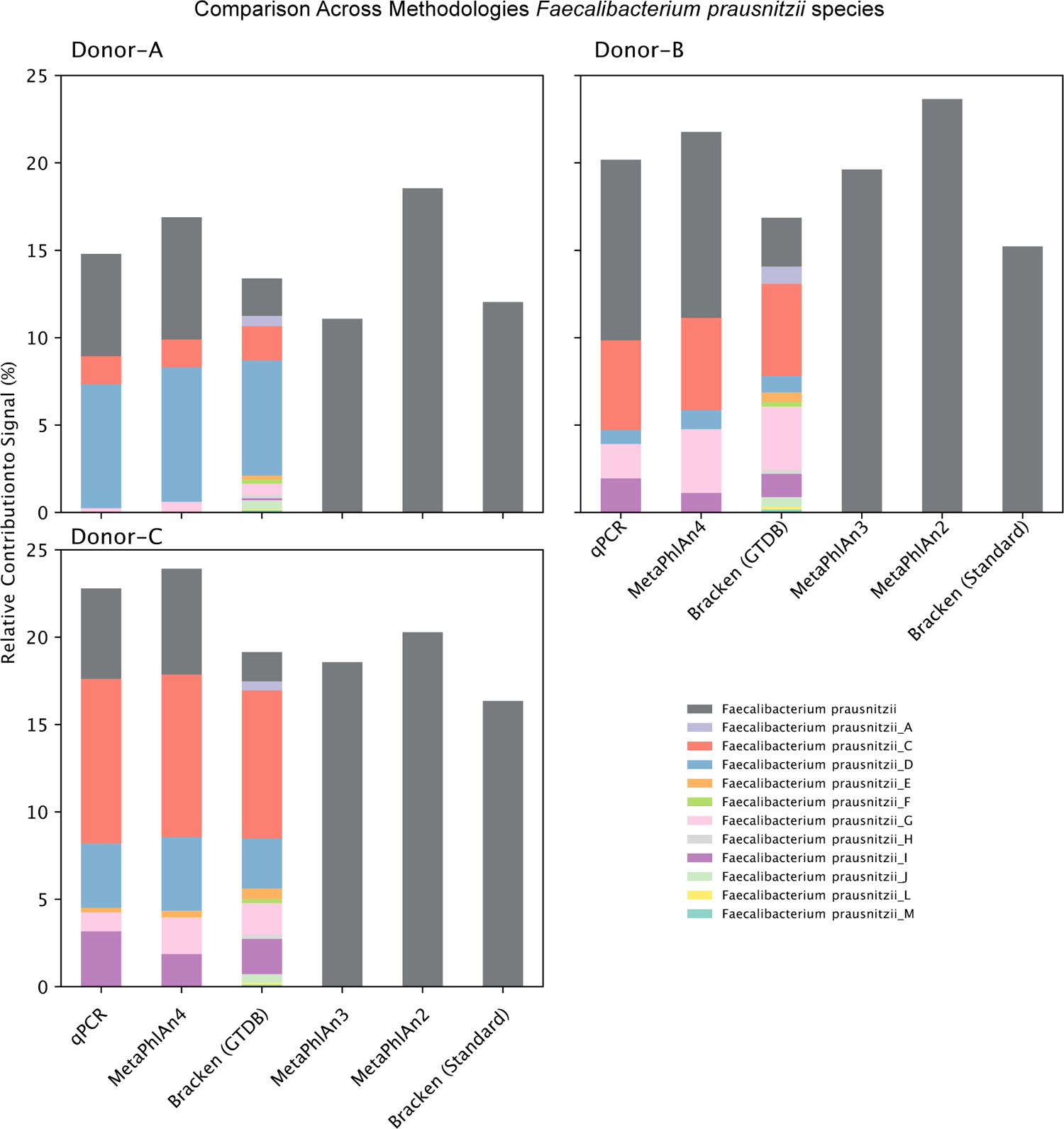
Comparison of relative contribution to signal for all three donors examining the 12 species complex of *Faecalibacterium prausnitzii*.

### 3.5 Limitations of detection - qPCR vs Bioinformatics

In examining the deep sequencing dataset, with the targeted subset of species, we identify 40 instances across all three (12-16 per sample) samples where taxa are detected by qPCR that cannot be definitively categorized by metagenomic approaches (Table 1 & 4). This is in contrast to database differences (as discussed above), but depending on the tool/database, either a low abundance microbe was not present in the output summary or recorded at such a low abundance that the signal is indistinguishable from background noise in the taxonomic assignment process. For a majority of these instances (40%), the number of detected copies is low (*<*10 copies per reaction). But nine taxa are present in samples with *>*100 copies per reaction, with two instances of taxa at 4,135 and 6,211 copies for *Streptococcus parasanguinis F* and *Bifidobacterium pseudocatenulatum*, respectively (approximately equivalent to *>*0.40% relative abundance for MetaPhlAn4 and Bracken-GTDB). When profiling the gut microbiome, microbes that fall below the detection threshold for metagenomic approaches can be derived from a number of locations along the digestive tract and correctly identifying and quantifying those microbes would impact interpretation of results. For example, using fecal samples to detect stomach/esophageal infections, monitor the engraftment of live therapeutics or probiotics in the upper gastrointestinal tract, or identify disease biomarkers.

Conversely, the metagenomic bioinformatic approaches identified 29 instances across all three samples (7-12 per sample) where taxa were not detected by qPCR. In each sample, MetaPhlAn2/3 shared exactly one taxon detected that was not identified by qPCR - *Ruminococcus E bromii* (Donor-A and -C) and *Hungatella A hathewayi A*. Both taxa represent species that were split using genome taxonomy which are lumped with the canonical species names in the MetaPhlAn2/3 databases. A majority of these instances were detected using Bracken with both the GTDB and Standard databases. As demonstrated with the artificial metagenomes, Bracken-based methods have issues with “read bleed” where closely related organisms recruit k-mers even when not present in the sample (Figure 3). This known issue is reflected in the higher “quantifiably reliable” cutoff value (0.04% versus 0.01%) empirically determined by Parks *et al*. (2021). Every instance detected by Bracken-GTDB is a species complex that has been split using the genome taxonomy approach. While Bracken-Standard did struggle with “split species groups”, the method also detected instances of species without this recent taxonomic update. However, no other method detected these organisms. The inaccuracies and limitations of the Standard database may play a significant role as the limited number of species become acceptable targets for more divergent k-mers. The ability to parse and identify likely causes for these discrepancies increases our confidence that the qPCR assays reflect biological reality and further highlight how bioinformatic methodological issues may confound microbiome associations.

**Figure 3.**
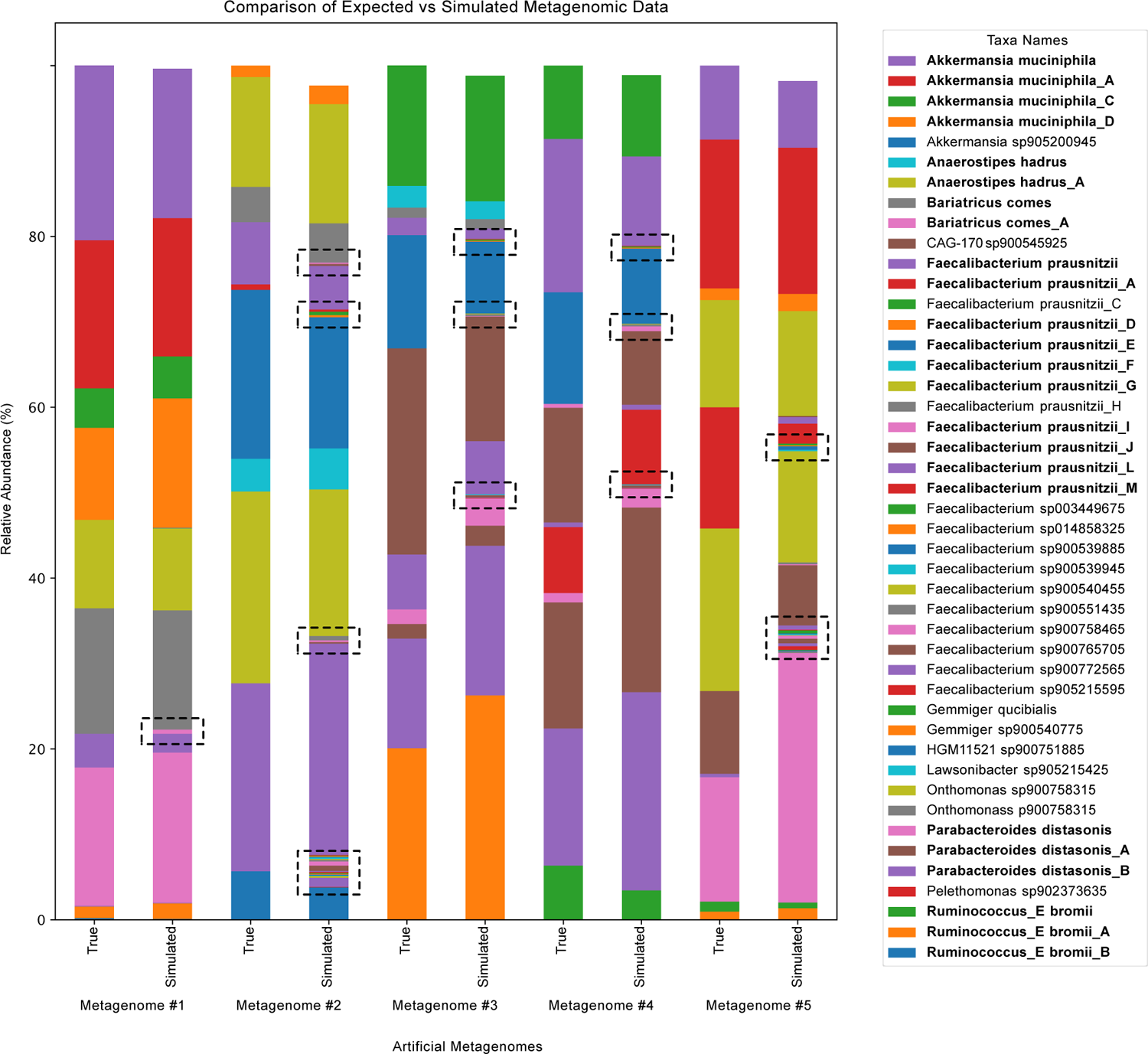
Five replicates of the artificial metagenomes with corresponding relative abundance results of Bracken-GTDB. The dotted boxed indicate taxa with visibly discernible abundance incorrectly detected by Bracken-GTDB. Note: In the figure legend, bold indicates taxa which were seeded into the artificial metagenomes.

### 3.6 Statistical outliers between methods

For every outlier taxa, if there is a disagreement between the qPCR and MetaPhlAn4 there is no disagreement with Bracken-GTDB and vice versa (Supplemental Data 4). Due to the way validation is performed for the qPCR assays, we do not expect either bioinformatic method to be consistently the benchmark of accuracy for the assay in a complex sample. Instead, especially due to the fluid nature of these taxonomic assignments and inconsistencies across databases (*e.g.*, *Anaerostipes hadrus* & *Anaerostipes hadrus A*; *F. prausnitzii*), the bioinformatic results act as guides to identify assays which are extreme outliers for both methods. There were no results in this dataset to suggest any of the qPCR assays produced extreme outlier information when anchored to these two methods. We do observe high variability and inconsistencies between qPCR and MetaPhlAn2/3 and Bracken-Standard. However, both MetaPhlAn2/3 are constructed on outdated reference databases/taxonomic structure and Bracken-Standard is limited by the size of its respective genome database.

### 3.7 The impact of “shallow” sequencing

“Shallow” sequencing is often a term used by companies that offer sequencing services to denote a type of sequencing that is more cost-effective per sample. In general, “shallow” sequencing typically produces about 10M metagenomic reads (5M paired-end reads) about 25*×* less than the “deep” sequencing performed for this dataset. When implementing “shallow” sequencing, as represented by random subsampling of the “deep” sequencing dataset, the number of taxa only detected by qPCR increased to 44 instances (Supplemental Data 2). This is important to highlight because it defines a key issue for planning research that requires microbiome profiling. “Shallow” sequencing can easily miss low abundance microbial signals, however additional sequencing depth and the associated cost does not resolve the underlying limit of detection issue for a vast majority of the missed taxa.

## 4 CONCLUDING REMARKS

While historically considered to be insufficient for large-scale microbiome profiling, we demonstrate here that with the correct infrastructure for *in silico* and *in vitro* validation qPCR molecular detection can provide accurate and quantitative microbiome profiles from complex natural samples. Our qPCR microbiome profiles are highly correlated with metagenomic bioinformatic approaches with added benefit of quantitative measures and the ability to detect exceedingly low abundance microorganisms. The ability to confidently detect microbes at low abundances may be a necessary requirement for research purposes to accurately profile a complex microbial community and can improve detection in samples with significant amounts of carryover DNA (*e.g.*, skin, oral, and urogenital microbiomes). The known underlying issues with metagenomic bioinformatic approaches can lead to high levels of uncertainty across samples and research groups. As demonstrated here, qPCR microbiome profiles offer a way to standardize detection and provide an avenue for reproducible and reliable microbiome data. The 110 qPCR assays discussed here function as a baseline proof-of-concept that can be further developed into several human microbiome panels capable of quantifying hundreds of species.

## ACKNOWLEDGMENTS

Thank you to the financial and commercialization support by the Alfred E. Mann Institute of Biomedical Engineering (now USC-AMI). Additional thanks to Drs. Jain, Schmid, Blanco, Lasch and Hong, Mr. Shiva, and Mr. Hutchinson and all the other AMI-USC staff for the support to this point.

