## Supplementary Information for "Benchmarking A Novel Quantitative PCR-based Microbiome Profiling Platform Against Sequencing-based Methods"

#### This PDF file includes:

Supplementary text
Figs. S1 to S2
Tables S1 to S2
Legends for Dataset S1 to S5
SI References

#### Other supplementary materials for this manuscript include the following:

Datasets S1 to S5

### Supporting Information Text

### Materials and Methods

**A. Assay Design.** Molecular assay *in silico* design and validation is performed by two Branchpoint Biosciences proprietary software packages, TaxaPrime™ and TaxaClear™, respectively. TaxaPrime™ determines a comprehensive list of all genomic regions in a target taxon (*i.e.*, a designated species group as defined by GTDB taxonomy (1)) that have properties required for a primer and/or probe whose quality is scored using a proprietary point system. Primer/probe compatibility is determined and all amplicons, including accompanying probe sites, are recorded. TaxaClear™ identifies and removes primers and probes with predicted cross-reactivity to non-target taxa. TaxaClear™ screens through two distinct phases. Phase one uses a database that consists of one representative for each species with known association with human hosts ( 4,200 species) plus the GRCCh38.p14 human reference genome. This phase screens for unintended targets in distantly related taxa and the human genome that would confound detection in a sample. Phase Two is semi-unique to each target taxon and consists of a database logically designed to comprise genomes of phylogenetically closely related species with a high probability (*i.e.*, overlapping body sites) to conflict with TaxaPrime™ results. This phase screens for potential cross-reactivity against near neighbors and improves specificity in complex samples.

**B. Deep Validation for 17 Targets.** To assess the effectiveness of TaxaPrime™ and TaxaClear™ in identifying species-specific primer pairs with compatible internal probe sites, qPCR assays were designed for 17 target species (Figure S2). For each target, 10 assays were selected for *in vitro* validation. Forward and reverse primers were ordered for all 170 assays (Integrated DNA Technologies, Inc., Iowa, USA) and each pair was assessed against genomic DNA from a diverse panel of 32 representative microorganisms observed in the human gut. Primer sets were assessed to confirm appropriate products were obtained from target organisms when present in the panel ($C_t$ <30) and that non-target organisms did not result in unintended cross-reactivity. Primer sets without any unexpected signal from non-target species ($C_t$ >35) were considered passing, whereas those resulting in any unexpected products from non-target species ($C_t$ <30) were considered failures. Instances of inefficient amplification from unintended species ($C_t$ between 30-35) were flagged for concern.

### Results

All 170 assays were assessed against a diverse panel of 32 representative microorganisms found in the human gut and scored according to their performance. A matrix summarizing the results of each assay is shown in Figure S2. Successful assays (green) amplified the intended product from the target species when present ($C_t$ >30) and exhibited no cross-reactivity with non-targets ($C_t$ >35). Assays that produced unintended signal ($C_t$ <30) from one or more non-target species were considered failures (red). Some assays produced one or more instances of only inefficient amplification from non-targets ($C_t$ 30-35) and were flagged for caution (yellow). Subsequent experiments (data not shown) demonstrated the addition of species-specific fluorogenic probes to primer sets flagged for caution successfully could distinguish target species from non-target species, but primer sets without any primer cross-reactivity were preferentially selected for subsequent validation within diverse stool communities.

### Discussion

TaxaPrime™ and TaxaClear™ successfully identified species-specific primers for all 17 targets. When initially assessed against a diverse panel of representative strains, 57% (97/170) successfully passed cross-reactivity validation, and at least 2 species-specific primer sets were identified for each target species. Furthermore, at least one of the first three primer sets assessed successfully passed cross-reactivity screening for 82% (14/17) of targets. The high success rate of assay design afforded by TaxaPrime™ and TaxaClear™ enabled the rapid and efficient expansion from 17 validated targets to the 110 assays assessed against stool communities within 2 months.

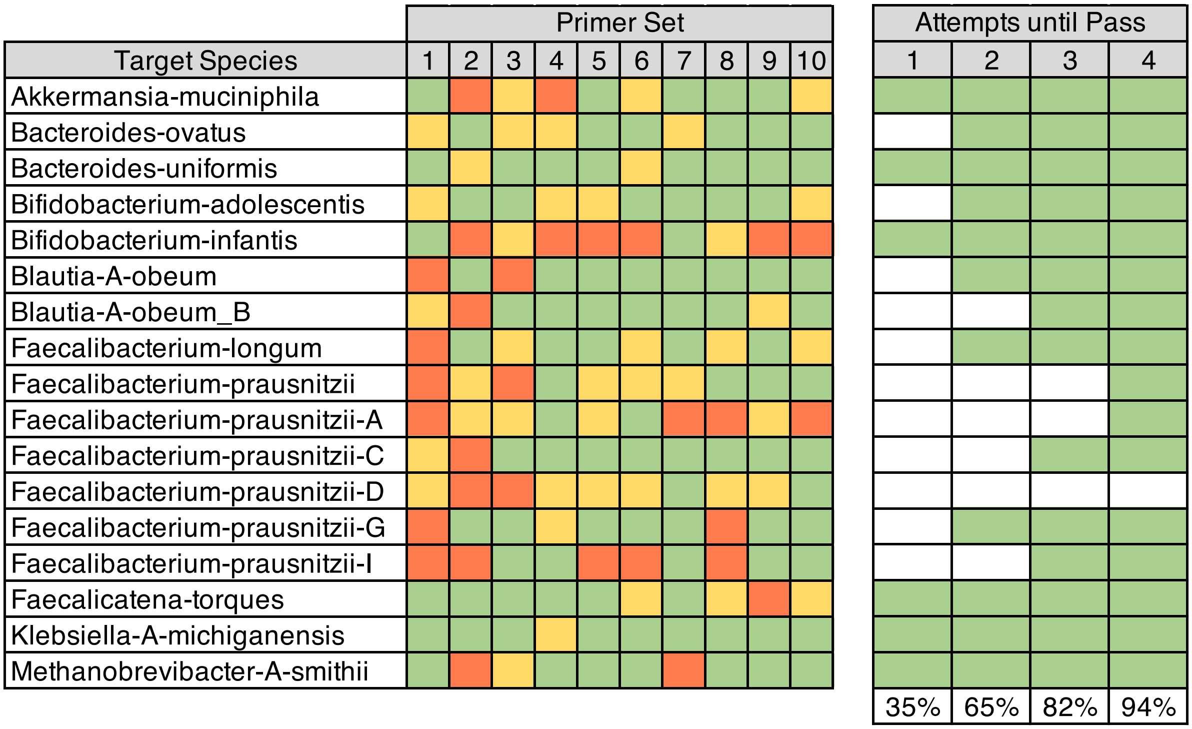


**Fig. S1.** Left panel: Successful assays (Ct >30; green) amplified the intended product from the target species and exhibited no cross-reactivity with non-targets ($C_t$ >35). Assays that produced unintended signal ($C_t$ <30; red) from one or more non-target species were considered failures. Some assays produced one or more instances of only inefficient amplification from non-targets ($C_t$ 30-35; yellow). Right panel: The number of sequential qPCR assays required during *in vitro* validation before identifying a successful assay. Columns 1-4 correspond to columns 1-4 in the right panel. Once a successful (green) assay is validated, remaining columns are set to success (green). The only target without a valid qPCR assays by the fourth column is *Faecalibacterium prausnitzii_D*, which required 7 *in vitro* tests before validation.

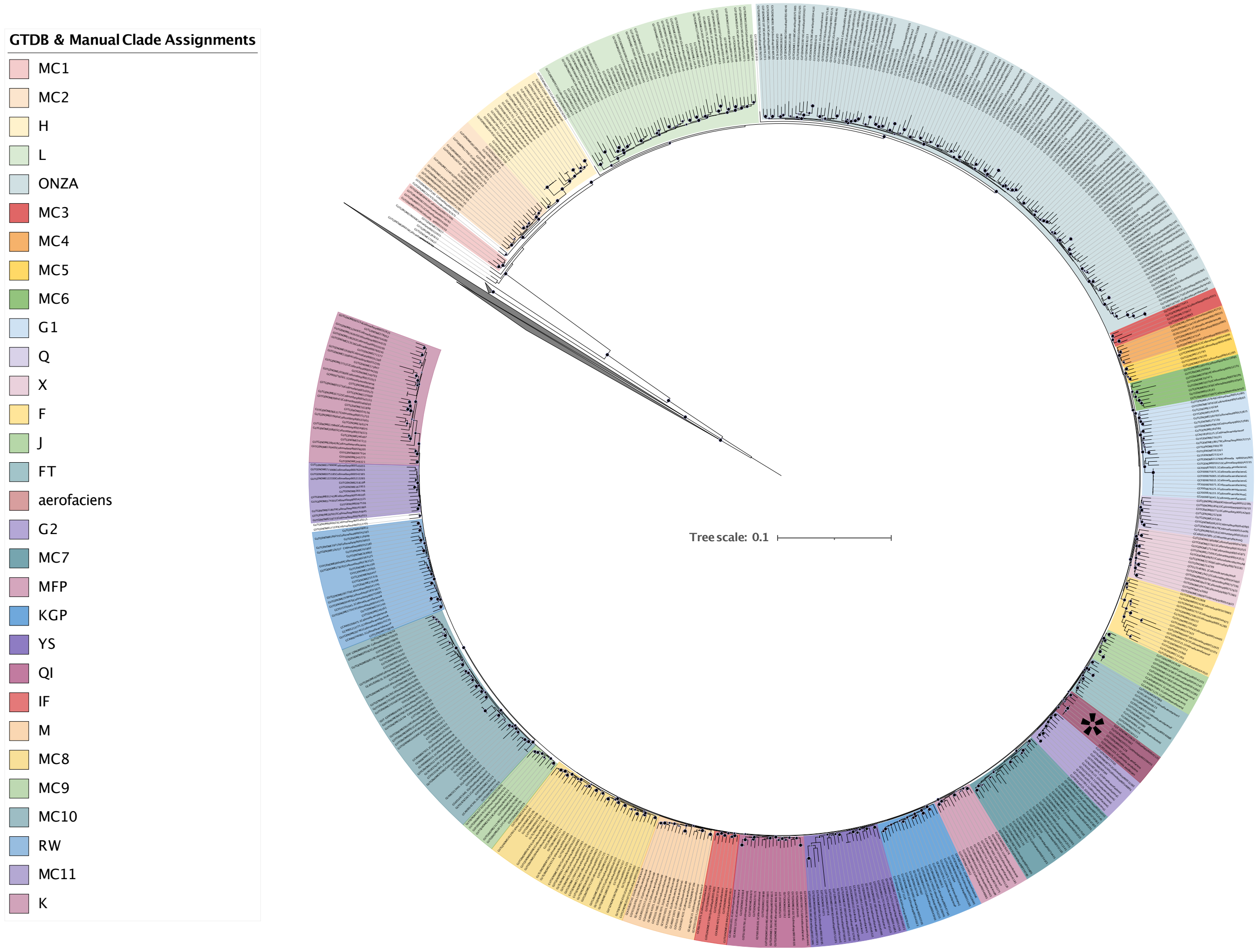


**Fig. S2.** Phylogenetic tree constructed using GToTree for 634 genomes. Color legend corresponds to GTDB clades and manual clades (MC). Phylogenetically cohesive units with aggregate multiple GTDB assignments are indicated (*e.g.*, clade ONZA contains GTDB assignments O, N, Z, and A. Manual clades contained genomes without or with few, inconsistent GTDB assignments. Despite extensive partitioning, viable qPCR assays were unachievable, however, one *Collinsella aerofaciens* encompassing set was possible. Black dots denote bootstrap values >0.75 and are scaled proportionally. The astericks denotes the clade that contains the *C. aerofaciens* type strain.

**Table S1. Fraction of reads assigned to a species using Kraken2/Bracken with the GTDB or Standard database**

| Sample | Total Reads (M read pairs) | GTDB (%) | Standard (%) |
|---|---|---|---|
| Donor-A | 132 | 81.46 | 42.37 |
| Donor-B | 146 | 83.71 | 50.21 |
| Donor-C | 116 | 83.71 | 51.47 |

**Table S2. Number of taxa report for each bioinformatic method above the "reliably quantifiable" threshold**

| Sample | MetaPhlAn4 | MetaPhlAn3 | MetaPhlAn2 | Bracken-GTDB | Bracken-Standard |
|---|---|---|---|---|---|
| Donor-A | 152 | 92 | 74 | 193 | 113 |
| Donor-B | 202 | 96 | 64 | 230 | 104 |
| Donor-C | 196 | 108 | 79 | 245 | 111 |

**SI Dataset S1** (**Supplemental-Data-1.xlsx**)

Raw data counts for compared methdods: qPCR genome equivalent copies, MetaPhlAn4 relative abundance, and read counts for MetaPhlAn3, MetaPhlAn2, Bracken-GTDB, and Bracken-Standard.

**SI Dataset S2** (**Supplemental-Data-2.xlsx**)

Summarized bioinformatic results as above or below the corresponding "reliably quantifiable" thresholds. Highlights taxa detected by qPCR or metagenomics. "Reliably quantifiable" defined as <0.4% for Bracken-based results and <0.1% for MetaPhlAn-based results.

**SI Dataset S3** (**Supplemental-Data-3.xlsx**)

Raw DADA2 combined taxonomy and relative abundance data.

**SI Dataset S4** (**Supplemental-Data-4.xlsx**)

Expanded comparative table for each sample showing outlier values for pairwise comparisons between methods.

**SI Dataset S5** (**Supplemental-Data-5.xlsx**)

GTDB Species, NCBI Genome Assembly, and relative abundance distributions used to construct the artificial metagenomes.
